# Population genomics highlight the vulnerability of coral-dwelling gobies to ecological losses due to climatic disturbances

**DOI:** 10.1101/2025.08.28.672959

**Authors:** Catheline Y.M. Froehlich, Renae L. Kirby, Samuel Greaves, Siobhan J. Heatwole, O. Selma Klanten, Martin Hing, Marian Y.L. Wong

## Abstract

Predicting species resilience to climate change generally focuses on species life history and ecology, overlooking genomic changes following disturbances. After consecutive cyclones and heatwaves, coral-dwelling *Gobiodon* gobies have experienced extreme population and group size reductions, and slower recovery rates than their coral hosts. To assess genomic contributions to this vulnerability, we analyzed multiple *Gobiodon* species for population structure (1) among three locations in western Oceania and (2) pre- and post-disturbances at the central location that underwent the same cyclones and heatwaves that reduced populations and group sizes. Species compared among locations all exhibited population structure. Post-disturbances, two species (*G. fuscoruber, G. rivulatus*) were extirpated, two species (*G. histrio, G. quinquestrigatus*) exhibited genetic bottlenecks, one species had no change to genetic diversity (*G. brochus*), and another species had consistently low diversity (*G. erythrospilus)*. Our results highlight that coral-dwelling gobies face compounding ecological and genomic losses from climatic disturbances, warranting targeted conservation.

## Introduction

Climate change is driving more frequent and intense disturbances, reshaping ecosystems at an alarming pace (AghaKouchak *et al*. 2018; Hughes *et al*. 2018a; Seidl *et al*. 2017). Most ecosystems face one or more disturbances, such as drought, floods, fires, storms, heatwaves, ocean acidification, increasing salinity, species invasions, and pollution (Bergstrom *et al*. 2021). This can lead to a variety of changes, like disease outbreaks, habitat degradation, behavioral modifications, altered species interactions, population declines, extirpations, and even ecosystem collapse (Sergio *et al*. 2018; Thakur *et al*. 2022; Turner 2010). To understand to what extent organisms are facing climate-related threats, it is crucial to study the impacts of climate change across multiple levels of biological organization (Attrill & Depledge 1997; Borges *et al*. 2021; Hourigan *et al*. 1988; Krishnan *et al*. 2013; Pinna *et al*. 2023). While ecological and physiological studies show that most species face mounting pressures due to climate change (Hill *et al*. 2021; Hughes *et al*. 2018b; Johnstone *et al*. 2016; Pinsky *et al*. 2022), understanding the extent of gene flow among populations has also been central to predicting their resilience and refining management strategies (Baptista *et al*. 2025; Cooke *et al*. 2016; Pauls *et al*. 2013; Woodall *et al*. 2011). By investigating the link between population structure and the potential for genetic bottlenecks, we can provide greater insights into species persistence with continued disturbances and guide effective management strategies (Jangjoo *et al*. 2016; Munguía-Vega *et al*. 2015; Ngeve *et al*. 2023). By combining insights from ecology, population connectivity, and genetic bottlenecks, we can better forecast species’ resilience and refine conservation strategies that aim to mitigate the impacts of climate change (Teng *et al*. 2025).

Reef fishes face major setbacks immediately following climatic disturbances, with coral-dwelling fishes being particularly vulnerable (Bellwood *et al*. 2012; Froehlich *et al*. 2021; Richardson *et al*. 2018). Our recent studies have shown that coral-dwelling gobies (genus *Gobiodon*) declined considerably after consecutive cyclones and bleaching events, as they suffered extreme population losses, and recovered slower than their coral hosts (Froehlich *et al*. 2021, 2023). Nine months after these consecutive disturbances, only a few adult and juvenile gobies were found. Although the community of coral hosts differed three years post-disturbances, only a few goby species switched hosts and the majority of corals were left uninhabited (Froehlich *et al*. 2023). When forcibly displaced—as can occur during cyclones—gobies overwhelmingly risked predation to return to their home coral rather than accepting a new host (Froehlich *et al*. 2022). Post-disturbance, these gobies no longer lived in groups (Froehlich *et al*. 2024), despite the well-documented benefits of group living for their survival and reproduction (Wong & Buston 2013). Together, these findings paint a troubling picture of severe and multifaceted changes to goby ecology and population dynamics following extreme climatic events. Such losses raise crucial questions about whether gobies possess long-term resilience to ongoing climate change.

Here, we combined two complementary approaches to explore the interplay between genetic connectivity and diversity of *Gobiodon* populations and declines in their populations from climatic disturbances. We asked two questions: 1) from a spatial perspective, is there consistent gene flow among different areas within their ranges that could facilitate recovery? And 2) from a disturbance perspective, can genetic bottlenecks be detected only three years after consecutive disturbances that triggered ecological and population declines? To address these questions, we sampled goby populations at three locations when in relatively healthy states: Papua New Guinea (Kimbe Bay), northern Great Barrier Reef (Lizard Island), and southern Great Barrier Reef (One Tree Island). We then revisited Lizard Island after the disturbances that resulted in extreme population and ecological (i.e. host types and sociality) losses to determine whether there were also losses in genetic diversity and signals of genetic bottlenecks. Together, these approaches revealed how ecological changes and mass mortality following extreme disturbances were coupled with population structure to result in extirpations and genetic bottlenecks for several *Gobiodon* species. Our study underscores the high vulnerability of coral-dwelling gobies to climate disturbances and need for additional protection.

## Methods

### Study Locations

Sampling was completed at three different locations in western Oceania (Fig. 1). The northern most location was Kimbe Bay, Papua New Guinea (PNG: -5.42896°, 150.09695°). We visited PNG from 25 Sep-3 Nov 2018 when the reef was relatively undisturbed. The central location was Lizard Island, northern Great Barrier Reef in Australia (LI; -14.687264°, 145.447039°). We visited LI from 30 Oct-5 Nov 2013 and 4 Feb-26 Feb 2014 when the reef was relatively undisturbed, and then from 15 Jan-5 Mar 2020 after the reef had been exposed to four consecutive years of extreme disturbances (category 4 cyclones Ita in 2014 and Nathan in 2015, and two mass bleaching events from extended marine heatwaves in 2016 and 2017). The southernmost location was One Tree Island, southern Great Barrier Reef in Australia (OTI; -23.506565°, 152.090954°). We visited OTI from 28 Jan-19 Feb 2019 when the reef was relatively undisturbed. The direct distance (point to point) between PNG and LI was ∼1140 km, between LI and OTI was ∼1200 km, and between PNG and OTI was ∼2000 km.

**Fig. 1.**
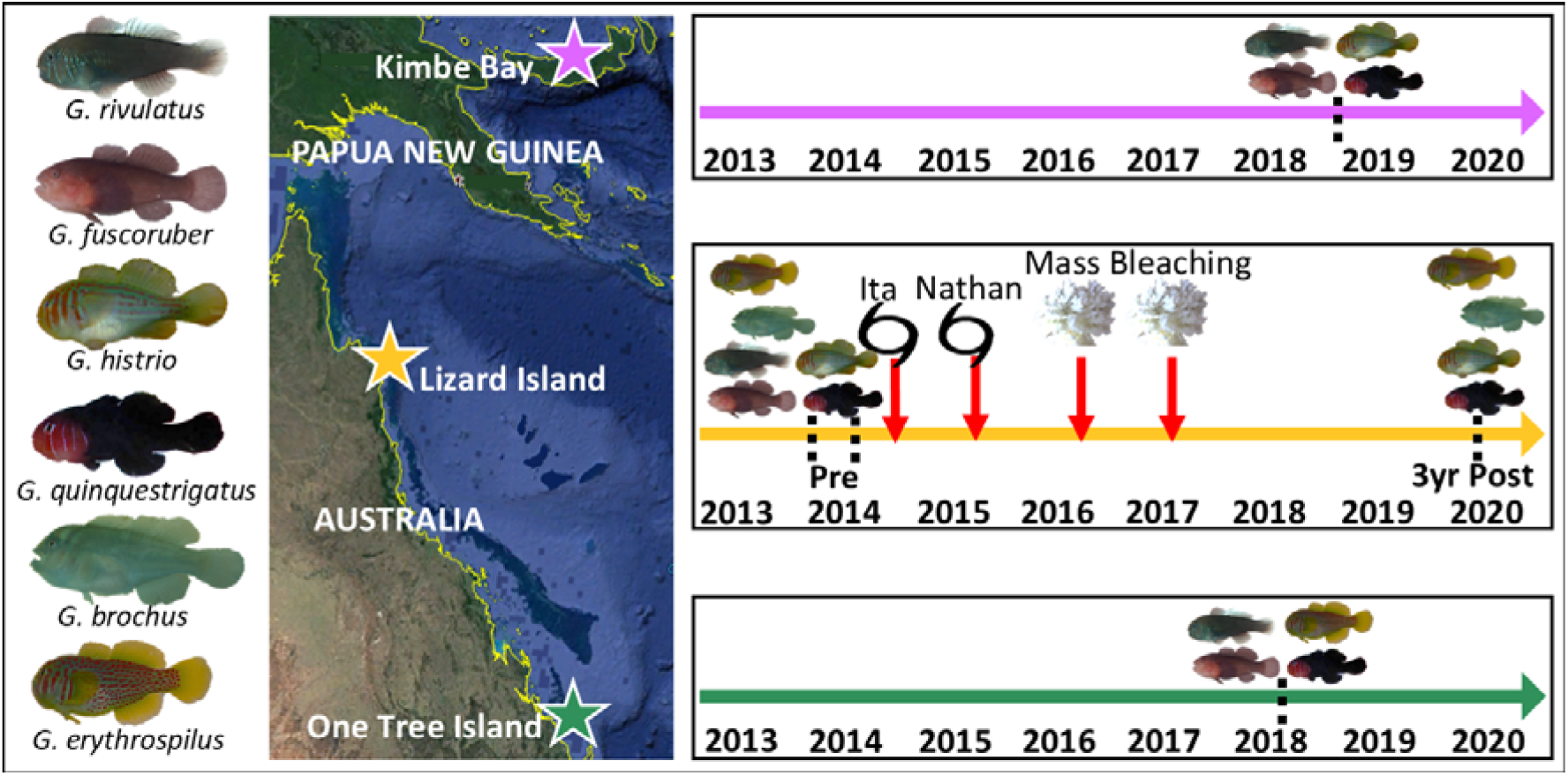
Three sampling locations with timelines of sampling events (dashed lines) and climatic disturbances (downward arrows; black spiral symbol for cyclones and white coral symbol for marine heatwaves that caused mass bleaching) for all species studied in western Oceania.

### Sample Collection

Tissue collection was completed on SCUBA. At each location, we searched for gobies within their *Acropora* coral hosts with the aid of an underwater flashlight. We selected goby species based on their abundance at all locations. When encountered, gobies were temporarily removed from the coral using an anesthetic solution (1:1:1 clove oil: seawater:70% ethanol) for anesthesia and a hand wafting motion around the coral to direct them into hand nets.

Gobies were identified to species level and a small fin clip was collected from their caudal fin, i.e. tail. Individuals were then returned to their coral hosts once they had recovered from the anesthesia. Collected tissues were stored in 80% ethanol.

### DNA Extraction and Sequencing

From Papua New Guinea, we sequenced 36 *G. fuscoruber*, 37 *G. histrio*, 43 *G. quinquestrigatus*, and 31 *G. rivulatus*. From Lizard Island pre-disturbances, we sequenced 47 *G. brochus*, 43 *G. erythrospilus*, 38 *G. fuscoruber*, 50 *G. histrio*, 53 *G. quinquestrigatus*, and 47 *G. rivulatus*. From Lizard Island post-disturbances, we sequenced 33 *G. brochus*, 33 *G. erythrospilus*, 40 *G. histrio*, and 41 *G. quinquestrigatus*. From One Tree Island, we sequenced 31 *G. fuscoruber*, 29 *G. quinquestrigatus*, and 29 *G. rivulatus*. We extracted DNA from the tissue at the University of Wollongong and Sydney Institute for Marine Science using the SV Genomic DNA Purification System (Promega) following the manufacturer’s protocol. We concentrated the DNA at 60°C in a vacuum centrifuge. We sent the samples to the Australian Genome Research Facility (AGRF) for library preparation and sequencing. Samples were prepared following the Peterson et al. (2012) protocol for double digest restriction-site associated digest (ddRAD) with restriction enzymes EcoRI and MseI, and size selected for a 60 bp region (350-410 bp fragments). The samples were sequenced on an Illumina NextSeq 500 (1x150 cycles mid-output) at AGRF.

### Data Analysis

Initial quality control and demultiplexing was performed by AGRF using STACKS v.2.3d software (Catchen *et al*. 2013, 2011). Sequences that had exact barcodes were retained and were then trimmed to 150bp. We then performed genotyping and Single Nucleotide Polymorphism (SNP) filtering using STACKS v2.54 (Rochette *et al*. 2019). Parameters used for denovo pipeline were as follows: six for the distance between stacks (-M), six for the distance between catalogue loci (-n) and three for the minimum stack depth of coverage (-m). These values were optimized following Rochette & Catchen (2017). Samples with less than 50% of loci called were excluded from analysis.

The data was divided into datasets for analyzing (1) differences among locations in relatively healthy states (PNG 2018, LI 2014, OTI 2019) for each species and (2) differences among pre- (2014) and post-disturbances (2020) (LI only) for each species. By using the “populations” component of STACKS, loci were exported by calling more than 80% of samples (-r) that were present in all populations (-p; 3 for among locations, except 2 for *G. histrio* which was only found at two locations; 2 for pre/post disturbances), had less than 0.7 observed heterozygosity (--max-obs-het), and had a minimum of 2 minor allele count (--min-mac). If loci were significantly out of Hardy-Weinberg Equilibrium (HWE) in two or more sampling locations, then they were removed. To properly calculate HWE, the Hardy-Weinberg (--hwe) option was used within populations and the p-values were then adjusted with the “BY” correction of the “p.adjust” function in R v3.6.3 (Rstudio V.1.4.1106) (R Core Team 2020; RStudio Team 2020) to account for False Discovery Rate. To avoid closely linked SNPs, the --write-single-snp option was used to export the final set of loci.

Population structure among the different locations (PNG, LI, OTI;) and pre and post-disturbances (LI only) were quantified for analysis of molecular variance (AMOVA) using Arlequin v3.5.2.2(Excoffier & Lischer 2010). For the AMOVA, 30,000 permutations were used and pairwise F_ST_ values were generated to assess hierarchical population structure among populations if the AMOVA was significant. The observed heterozygosity (H_0_), expected heterozygozity (H_e_), and inbreeding coefficient (F_IS_) were calculated as genetic diversity statistics using the “basic.stats” function in the R package hierfstat (Goudet 2005). To determine whether there was asymmetric genetic migration, we used the divMigrate function (Sundqvist *et al*. 2016) from the diveRsity package (Keenan *et al*. 2013) in R, using the “Nm” statistic to estimate number of migrants, and 1000 bootstrap samples when calculating significance.

To identify patterns of admixture in the populations with each analysis, STRUCTURE v.2.3.4 (Pritchard *et al*. 2000) was used. In each analysis, we included 100,000 iterations for burn-in and 500,000 iterations of Markov-chain Monte Carlo (MCMC), and for each K value (1-9) we used 9-25 independent runs per K depending on the species to limit the number of minor modes created. By applying the △*K* Evanno method (Evanno *et al*. 2005), the most likely K value was determined through STRUCTURE HARVESTER (Earl & vonHoldt 2012). The Cluster Markov Packager Across Ks (CLUMPAK) online tool (Kopelman *et al*. 2015) was used to summarize the results, and we produced the STRUCTURE graphs in R. The R package adegenet v1.7-16 (Jombart 2008; Jombart & Ahmed 2011) was used to conduct a Discriminant Analysis of Principal Components (DAPC). Loci were removed if there were more than 20% missing values, and individuals were removed if there were more than 25% missing values. For the remaining data, the missing values were imputed using the mean allele frequency for each locus calculated by the “tab” function. The “find.clusters” function was used to find the number of *K* values within each analysis regardless of a priori population groups sampled by using a combination of Bayesian Information Criterion (BIC) to identify the most likely K value, as well as a k-means approach to identify genetic clusters. The “xvalDapc” function was used to cross-validate results and identify the number of discriminant functions and principal components to retain for DAPC for each species per analysis.

## Results

### Population Structure Among Locations

After quality control and SNP calling, 101 individuals from *Gobiodon fuscoruber*, 103 *G. rivulatus*, 99 *G. quinquestrigatus*, and 75 *G. histrio* were retained with 5751, 6345, 6527, and 10006 loci respectively passing filtering parameters for the analysis among locations. Each species examined had distinct population structure across the locations where it was observed. However, the degree of clustering, heterozygosity and direction of migration asymmetry among locations varied by species.

*Gobiodon fuscoruber* exhibited the most distinct population structure among locations (AMOVA: F_ST_ = 0.14, P < 0.0001, see Suppl. Tabs. for all statistical outputs) compared to all other species (see results below). There was some shared ancestry between LI (central population) and PNG (northern population; pairwise F_ST_ = 0.12) as well as LI and OTI (southern population; pairwise F_ST_ = 0.11; Fig. 2A&B). However, PNG and OTI were highly distinct from each other with little shared ancestry (pairwise F_ST_ = 0.22). Regardless of population designation, three clusters were identified (⍰*K* = 3, 9 STRUCTURE HARVESTER repetitions per K, no minor modes; Suppl. Fig. 1). The smallest cluster included all PNG individuals whereas OTI and LI each had individuals with ancestry dispersed among the three clusters (Fig. 2B). The inbreeding coefficient (F_IS_) was high at LI, moderate at OTI, and low at PNG (Table 1). Heterozygosity values for both observed and expected were between 0.05 and 0.09 and observed (H_o_) was only higher than expected (H_e_) at PNG (Table 1). There was asymmetry in the migration rates from PNG to LI (100%) and PNG to OTI (61%; Fig. 2C), yet there was no asymmetry in migration between LI and OTI.

**Table 1.** Genetic diversity measures for each population of *Gobiodon* species studied. To investigate inbreeding coefficients, we used the range suggested by (Hartl 1999) in which 0-0.05 is low, 0.05-0.15 is moderate, 0.16-0.24 is high, and ≥0.25 is very high.

| Species | Population | Inbreeding Coefficient $F_{IS}$ | Observed Heterozygosity $H_o$ | Expected Heterozygosity $H_e$ |
| --- | --- | --- | --- | --- |
| *G. fuscoruber* | Papua New Guinea | 0.048 | 0.06 | 0.05 |
|  | Lizard Island | 0.174 | 0.07 | 0.09 |
|  | One Tree Island | 0.106 | 0.06 | 0.07 |
| *G. histrio* | Papua New Guinea | 0.125 | 0.08 | 0.09 |
|  | Lizard Island | 0.134 | 0.09 | 0.11 |
|  | One Tree Island | NA | NA | NA |
| *G. quinquestrigatus* | Papua New Guinea | 0.195 | 0.06 | 0.07 |
|  | Lizard Island | 0.156 | 0.05 | 0.06 |
|  | One Tree Island | 0.088 | 0.06 | 0.06 |
| *G. rivulatus* | Papua New Guinea | 0.137 | 0.07 | 0.08 |
|  | Lizard Island | 0.128 | 0.06 | 0.07 |
|  | One Tree Island | 0.061 | 0.04 | 0.04 |

**Figure 2.**
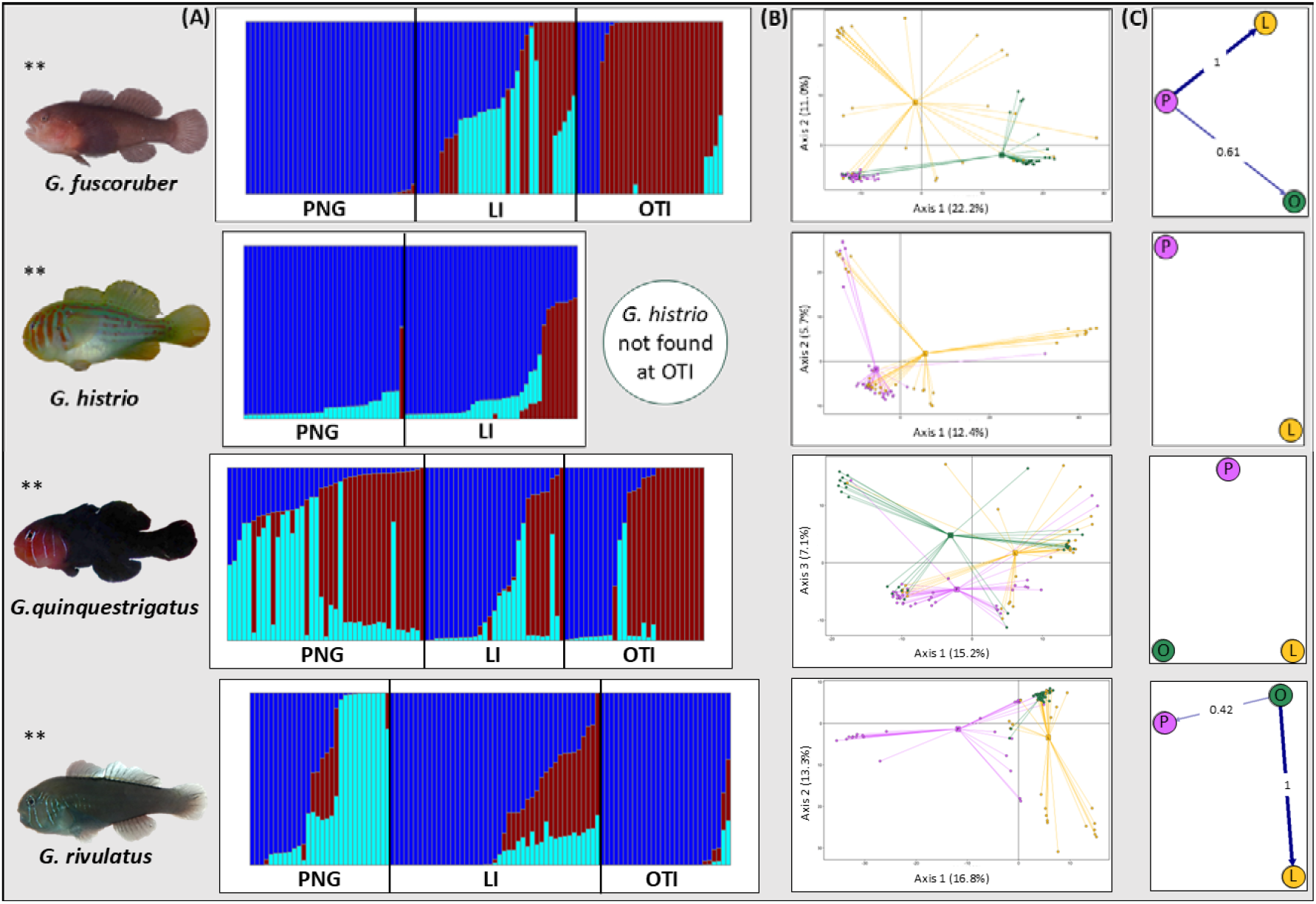
(A) Bayesian clustering algorithm STRUCTURE v2.3.4 with color representing different clusters following (Pritchard *et al*. 2000) and (B) Principal Components Analysis of population differentiation and (C) migration rates filtered for asymmetries for 3 sampled locations (purple = PNG, orange = LI, green = OTI) of each *Gobiodon* species. PNG = Papua New Guinea, LI = Lizard Island, OTI = One Tree Island. ** means p < 0.0001.

We found *Gobiodon histrio* at only two of the three sampling locations (PNG and LI). *G. histrio* had distinct population structure among the two locations (AMOVA: F_ST_ = 0.03, P < 0.0001), yet it had the least structure compared to other species. There were several individuals with shared ancestry among locations (Fig. 2A&B). Regardless of population designation, three clusters were identified depending on the analysis used (⍰K = 3, 25 STRUCTURE HARVESTER repetitions per K, no minor modes; Suppl. Fig. 1). However, no cluster fully encapsulated the population from a single location (Fig. 2B). The F_IS_ was moderate for both locations and slightly higher at LI (Table 1). Heterozygosity values for both H_o_ and H_e_ were between 0.08 and 0.11 with H_o_ being lower than H_e_ at both locations (Table 1). There was no asymmetry in the migration rates among locations (Fig. 2C).

*Gobiodon quinquestrigatus* had distinct population structure among the three locations (AMOVA: F_ST_ = 0.05, P < 0.0001), albeit with mixed ancestry present in all locations (pairwise F_ST_ values: 0.06 for PNG-LI, 0.03 for LI-OTI, 0.05 for PNG-OTI; Fig. 2A&B). Regardless of population designation, three clusters were identified (⍰*K* = 3, 25 STRUCTURE HARVESTER repetitions per K, 2 minor modes; Suppl. Fig. 1&2). However, no cluster fully encapsulated the population from a single location (Fig. 2B). The F_IS_ was high at PNG, high at LI, and moderate at OTI (Table 1). Heterozygosity values for both H_o_ and H_e_ were between 0.05 and 0.07 and H_o_ was lower or the same as H_e_ at all locations (Table 1). There was no asymmetry in the migration rates among locations (Fig. 2C).

*Gobiodon rivulatus* had distinct population structure among the three locations (AMOVA: F_ST_ = 0.09, P < 0.0001). There was shared ancestry between LI and OTI (pairwise F_ST_ = 0.04) and some between LI and PNG (pairwise F_ST_ = 0.10), but less between PNG and OTI (pairwise F_ST_ = 0.13; Fig. 2A&B). Regardless of population designation, three clusters were identified (⍰K = 3, 25 STRUCTURE HARVESTER repetitions per K, 2 minor modes; Suppl. Fig. 1&3). A single cluster encapsulated the PNG population and some individuals from LI and OTI, which were both dispersed among several clusters (Fig. 2B). The F_IS_ was moderate at all sites although it was two times higher at PNG and LI than OTI (Table 1). Heterozygosity values for both H_o_ and H_e_ were between 0.05 and 0.08 and H_o_ was lower or the same as H_e_ at all locations (Table 1). There is asymmetry in the migration rates from OTI to PNG (42%) and OTI to LI (100%, Fig. 2C), but no asymmetry between PNG and LI.

### Pre- versus Post-Disturbances at Lizard Island

*Gobiodon fuscoruber* and *G. rivulatus* were locally extirpated from Lizard Island following the climatic disturbances. For the pre- and post-disturbances subsets from Lizard Island, we retained 62 *G. brochus*, 70 *G. histrio*, 59 *G. quinquestrigatus*, and 62 *G. erythrospilus* individuals with 9724, 9391, 7656, and 10760 loci, respectively. Two out of the four species had distinct structure between pre- and post-disturbances and the degree of heterozygosity varied among species.

*Gobiodon brochus* had no distinct structure pre- versus post-disturbances (AMOVA: F_ST_ = 0.0007, P = 0.87). Regardless of population designation, four clusters were identified (⍰*K* = 4, 10 STRUCTURE HARVESTER repetitions per K, 0 minor modes; Suppl. Fig. 4). There was a loss of a single cluster made up of 2 individuals from pre- to post-disturbances (Fig. 3A&B). The F_IS_ was moderate at both time points (Table 2). Heterozygosity values for both H_o_ and H_e_ were between 0.14 and 0.17 and H_o_ was lower than H_e_ at both time points (Table 2). There was no asymmetry in the migration rates among sampling periods (Fig. 3C).

**Table 2.** Genetic diversity measures for each *Gobiodon* species studied pre- and post- disturbances at Lizard Island, northern Great Barrier Reef.

| Species | Population | Inbreeding Coefficient $F_{is}$ | Observed Heterozygosity $H_o$ | Expected Heterozygosity $H_e$ |
| --- | --- | --- | --- | --- |
| *G. brochus* | Pre-disturbances | 0.134 | 0.14 | 0.17 |
|  | Post-disturbances | 0.127 | 0.14 | 0.17 |
| *G. erythrospilus* | Pre-disturbances | 0.130 | 0.13 | 0.16 |
|  | Post-disturbances | 0.131 | 0.13 | 0.15 |
| *G. fuscoruber* | Pre-disturbances | 0.174 | 0.07 | 0.09 |
|  | Post-disturbances | extirpated | extirpated | extirpated |
| *G. histrio* | Pre-disturbances | 0.075 | 0.12 | 0.13 |
|  | Post-disturbances | 0.108 | 0.08 | 0.10 |
| *G. quinquestrigatus* | Pre-disturbances | 0.183 | 0.08 | 0.10 |
|  | Post-disturbances | 0.262 | 0.06 | 0.10 |
| *G. rivulatus* | Pre-disturbances | 0.128 | 0.06 | 0.07 |
|  | Post-disturbances | extirpated | extirpated | extirpated |

**Figure 3.**
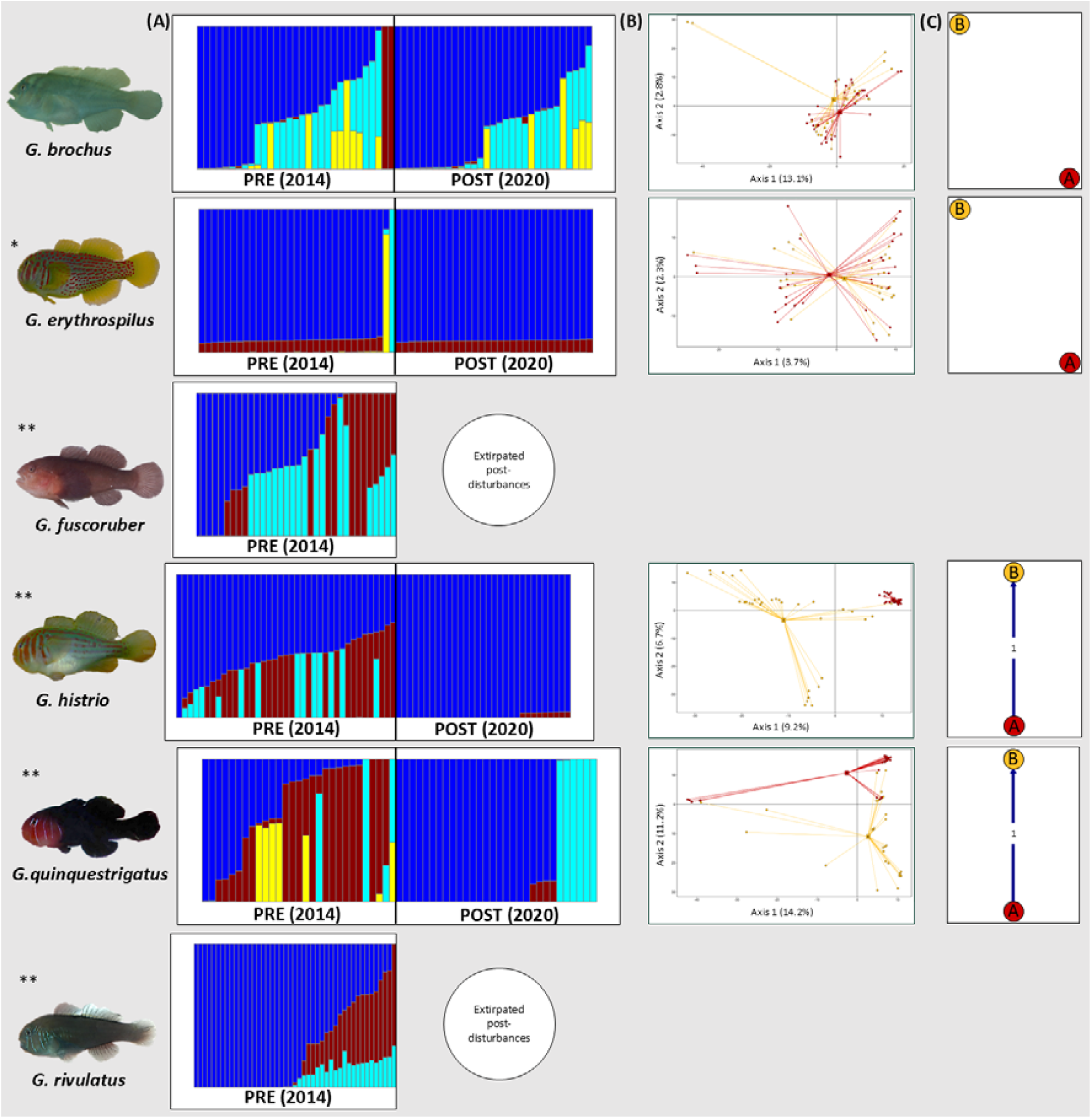
(A) Bayesian clustering algorithm STRUCTURE v2.3.4 with color representing different clusters (Pritchard *et al*. 2000) and (B) Principal Components Analysis showing population differentiation and (C) asymmetric gene flow migration for pre- (PRE = orange) and post-disturbances (POST = dark red) at Lizard Island of each *Gobiodon* species. ** means p < 0.0001, * means p < 0.02.

*Gobiodon erythrospilus* had barely significant population structure pre- and post-disturbances (AMOVA: F_ST_ = 0.005, P = 0.018). However, only one cluster was identified regardless of population designation (Suppl. Fig. 4). There were two clusters that disappeared from pre-to post-disturbances, although those clusters had only been found in two individuals pre-disturbances. The rest of the individuals exhibited the same ancestry at both time periods (Fig. 3A&B). The F_IS_ was moderate at both time points (Table 2). Heterozygosity values for both H_o_ and H_e_ were between 0.13 and 0.16 and H_o_ was lower than H_e_ at both time points (Table 2). There was no asymmetry in the migration rates between either time points (Fig. 3C).

*Gobiodon histrio* had distinct population structure pre- and post-disturbances (AMOVA: F_ST_ = 0.08, P < 0.0001). The population structure homogenized post-disturbances, with the loss of individuals with partial ancestry from another population (Fig. 3A&B). Regardless of population designation, three clusters were identified (⍰*K* = 3, 10 STRUCTURE HARVESTER repetitions per K, 0 minor modes; Suppl. Fig. 4). The F_IS_ was moderate at both time points but was 44% higher post-disturbances than pre-disturbances (Table 2).

Heterozygosity values for both H_o_ and H_e_ were between 0.08 and 0.13 and H_o_ was lower than H_e_ at both time points (Table 2). Although there was asymmetric migration identified from post-disturbances to pre-disturbances, the context instead implies the opposite direction (Sundqvist *et al*. 2016) with a founder effect due to the disturbances (Fig. 3C).

*Gobiodon quinquestrigatus* had distinct population structure pre- and post-disturbances (AMOVA: F_ST_ = 0.08, P < 0.0001). This species similarly homogenized post-disturbance, with almost all individuals exhibiting full ancestry from only 2 populations (Fig. 3A&B). Regardless of population designation, four clusters were identified (⍰*K* = 4, 10 STRUCTURE HARVESTER repetitions per K, 0 minor modes; Suppl. Fig. 4). Individuals pre-disturbances were found in each of the four clusters whereas one cluster was lost post-disturbances. Most individuals post-disturbances were from one of two clusters (dark blue or cian) with a select few exhibiting also third cluster (red with dark blue) (Fig. 3B). The F_IS_ was high pre-disturbances and 43% higher post-disturbances sitting at a very high value (0.26), which was the highest in the whole study (Table 2). Heterozygosity values for both H_o_ and H_e_ were between 0.06 and 0.10 and H_o_ was lower than H_e_ at both time points (Table 2). Although there was asymmetric migration identified from post-disturbances to pre-disturbances, the context instead implies the opposite direction (Sundqvist *et al*. 2016) with a founder effect due to the disturbances (Fig. 3C).

## Discussion

We used a genomic approach to quantify population structure and the potential for genetic bottlenecks resulting from the severe declines in *Gobiodon* populations and changes in their social structures and host-use patterns following repeated cyclones and heatwave-induced bleaching events (Froehlich *et al*. 2021, 2023, 2024). While all species showed some degree of population structure across the study sites, they differed in genetic diversity and directional gene flow. For example, *G. fuscoruber* exhibited migration patterns from north to south, the opposite was found for *G. rivulatus*, and *G. histrio* and *G. quinquestrigatus* had no clear pattern. Following disturbances at the central location, *G. fuscoruber* and *G. rivulatus* were extirpated. We detected genetic bottlenecks in *G. quinquestrigatus* and *G. histrio*. Our study highlights that population structure and genetic bottlenecks increases the vulnerability of these fishes to population collapse during climatic disturbances, alongside other ecological losses observed for theses fishes. Given these results, we recommend expanding conservation efforts to include more *Gobiodon* species, which are facing multiple climate-driven threats.

Population structure was evident for each *Gobiodon* species across all locations in relatively healthy states, suggesting these populations may be especially vulnerable to climatic disturbances. Most species exhibited low to moderate levels of genetic isolation, as indicated by F_ST_ values (0.03–0.13), while *G. fuscoruber* showed moderate to high F_ST_ values (0.11– 0.22) (Hall 2022). Although the distances between study sites (1000–2000 km) are typically below thresholds for genetic isolation at sea (Cooke *et al*. 2016), *G. fuscoruber*’s minimum F_ST_ of 0.15 (Frankham *et al*. 2002) indicates that its populations are indeed genetically isolated. Such populations may be either recruitment-limited or post-recruitment regulated (Jones *et al*. 2009). High self-recruitment rates are observed in many reef fishes with short larval durations (D’Aloia *et al*. 2018; Jones *et al*. 1999, 2005; Saenz-Agudelo *et al*. 2012; Selwyn *et al*. 2016; Swearer *et al*. 2002), like the coral-dwelling goby *Paragobiodon xanthosoma* (Rueger *et al*. 2021b). For *Gobiodon*, the pelagic larval duration averages around 19.6 days (Pereira *et al*. 2015), which suggests that they are regulated by post-recruitment and are not recruitment-limited under normal conditions, which reduces the risks of extirpation but increases recovery time (Jones *et al*. 2009). Nonetheless, two species (*G. fuscoruber* and *G. rivulatus*) experienced local extirpation in the disturbed location. Following these severe events, which dramatically reduced adult populations, *Gobiodon* may have become recruitment-limited as well. Inbreeding levels further highlight species-specific vulnerabilities. Inbreeding ranged from low to high (Hartl 1999) at different locations depending on the species (e.g. *G. fuscoruber, G. quinquestrigatus*), or moderate throughout (*G. rivulatus*). At the central location (Lizard Island, inbreeding was moderate in *G. histrio* and *G. rivulatus* and high in *G. fuscoruber* and *G. quinquestrigatus*. Such inbreeding patterns are unsurprising since *Gobiodon*, like other habitat specialists, tend to settle near natal reefs with siblings (Buston *et al*. 2009; Jones *et al*. 2005). However, limited genetic diversity may occur from such settlement behavior, therefore increasing their susceptible to extreme disturbances (Jones *et al*. 2009). Accordingly, low to high population structure coupled with moderate to high inbreeding patterns for all species likely contribute to their slow recovery rates.

Ocean currents play crucial roles in larval dispersal and the recovery potential of marine species (Gouvêa *et al*. 2023; Snead *et al*. 2023; Wilson *et al*. 2016), but they do not fully explain the patterns of directional genetic migration in *Gobiodon* species. Based on prevailing ocean currents in western Oceania (Ridgway & Hill 2009), the central populations in our study (Lizard Island) would be expected to act as a source for the northern populations (Papua New Guinea). Additionally, the presence of competing currents between the southern and central populations (East Australia and Hirri Currents; Ridgway and Hill 2009) further complicates the predictions of source-sink dynamics between these locations. Of the four species studied, only two species (*G. fuscoruber* and *G. rivulatus*) exhibited directional genetic migration, and both were locally extirpated from the disturbed location. Interestingly, their migration patterns were in opposite directions: *G. fuscoruber* showed directional genetic migration from the northern population to the central and southern populations, whereas *G. rivulatus* exhibited the opposite. It is possible that we did not sample intermediate populations found between the study locations, which may help explain the contrasting patterns among species. However, ocean currents alone may not explain the overall pattern of gene flow in all *Gobiodon* species, especially given that the central population would be expected to have sourced both the northern and southern locations. However, ocean currents are affected by climate change, which may alter patterns of larval dispersal (Wilson *et al*. 2016). Although the dispersal and genetic structure of many marine fishes is often driven by ocean currents (Baltazar-Soares *et al*. 2014; Munguia-Vega *et al*. 2017; Snead *et al*. 2023), not all tropical species exhibit such patterns (Shulman & Bermingham 1995). Since two *Gobiodon* species exhibited contrasting patterns despite their overlapping ranges, their genetic migration is likely influenced by other ecological factors, like their sociality (Froehlich *et al*. 2024; Rueger *et al*. 2021a) and the distribution of their preferred hosts (Froehlich *et al*. 2023). Overall, our findings indicate that ocean currents do not always influence population structure of all species within a single genus, which has flow-on implications for their resilience to climatic disturbances.

When investigating whether climatic disturbances at the central location affected the genetic diversity, not all species responded in the same manner. There were few changes to the population structure of *Gobiodon brochus* and *G. erythrospilus* following the disturbances. The latter species had little genetic diversity at both time points, which aligns with its vulnerable status (Larson 2020). Other aquatic taxa that underwent mass mortality following storms and heatwaves had little changes to their genetic diversity, such as the submerged aquatic plant *Vallisneria americana* and the red gorgonian *Paramuricea clavata* (Ngeve *et al*. 2023; Pilczynska *et al*. 2016). For the alpine butterfly *Parnassius smintheus*, high connectivity pre-disturbances resulted in little change to their genetic diversity after mass mortality (Jangjoo *et al*. 2016). However, not all species are resilient. Two species in our study, *G. fuscoruber* and *G. rivulatus*, were extirpated. The other two species, *G. histrio* and *G. quinquestrigatus*, showed a clear loss of genetic diversity, increased inbreeding, and marked directional genetic migration post-disturbance. In both species, post-disturbance populations were more genetically uniform (i.e. less diverse), and at least one ancestral cluster that was initially present pre-disturbances disappeared entirely. There were characteristics similar to a founder effects (Sundqvist *et al*. 2016) whereby a few individuals pre-disturbances gave rise to the post-disturbance populations. This was evident from the reduction in the number of ancestral clusters, the increase in assignment of individuals to a single cluster, and the directional genetic migration observed. Broader patterns in marine ecosystems show similar results in which population structure leads to lower genetic diversity post-disturbances and the potential for genetic bottlenecks (Noreen *et al*. 2009; Selkoe *et al*. 2016). Similarly, seaweed populations underwent up to 80-100% loss in genetic diversity following marine heatwaves (Gurgel *et al*. 2020). Taken together, our results suggest that genetic bottlenecks pose a real threat to several *Gobiodon* species. Conservation strategies aimed at protecting coral reef ecosystems should include coral-dwelling fishes and incorporate species-specific management plans at a population level. As climatic disturbances continue to increase in frequency and severity, genetic bottlenecks may become even more common, placing many more reef-associated species at risk.

When we look at the genetic, life history, and ecological outcomes of climate change on *Gobiodon* species, a clearer picture emerges as to why some species recover more slowly than others. For example, the two species that were extirpated post-disturbances (*G. fuscoruber* and *G. rivulatus*) had some of the highest F_ST_ values among populations. Pre-disturbances, both species lived primarily in groups rather than pairs, unlike other goby species in this study (Froehlich *et al*. 2024; Hing *et al*. 2019). Post-disturbances, the absence of established social groups for *G. fuscoruber* and *G. rivulatus* likely reduced successful settlement, compounded by high inbreeding rates and population structure. All *Gobiodon* species—regardless of whether they typically lived in pairs or groups before the disturbances—were mostly found living alone or in pairs post-disturbances (Froehlich *et al*. 2024), likely reducing their reproductive success. Additionally, a species’ growth potential also shapes resilience. For example, *G. rivulatus* is one of the smallest species in the genus (Hing *et al*. 2018, 2019), making it more vulnerable to predation, especially since smaller individuals are more frequently targeted post-disturbance (Brandl *et al*. 2018; Goatley & Bellwood 2016). Since *Gobiodon* species fully depend on live coral, how well their preferred coral hosts recover can directly affect their own recovery. *Gobiodon* species that were extirpated post-disturbances (*G. fuscoruber* and *G. rivulatus*) primarily inhabited corals that became rare post-disturbances (*Acropora millepora* and *A. gemmifera*, respectively) (Froehlich *et al*. 2021, 2023). However, *Gobiodon* species that persisted either were able to change the coral host they inhabited or benefited from their preferred host recovering quickly (Froehlich *et al*. 2023). Yet even species that survived were slower to recover than their coral hosts and several exhibited genetic bottlenecks, which further reduces their ability to respond to future disturbances. These compounding pressures highlight the urgent need for conservation efforts that specifically target coral-dwelling gobies as they play important roles for coral health and resilience to bleaching.

### Animal Ethics and Permits

The study was approved under animal ethics protocols AE1404 and AE1725 at the University of Wollongong. This research was conducted under research permits issued by the Great Barrier Reef Marine Park Authority (G13/36197.1, G15/37533.1, and G18/41020.1) and Papua New Guinea Research Visa Permit AA654347.

## Acknowledgments

We pay respect to the traditional owners past and present of the Kilu and Tamare communities of Kimbe Bay in Papua New Guinea, of the Dingaal Aboriginal Traditional Owners of Lizard Island in Australia, and Gurang, Gooreng Gooreng, Bailai and Bunda Traditional Owners of One Tree Island in Australia. We greatly appreciate our field helpers: Kylie Brown, Karen Hing, Theresa Rueger, Rituraj Sharma, Jemma Okapie Smith, Nelson and Jerry Sikatura. We are thankful to Mark Dowton and Tracey Gibson for guidance regarding laboratory practices. We are grateful to the facilities and management of field stations at the Mahonia Na Dari Research and Conservation Centre of Kimbe Bay, Lizard Island Research Station, and One Tree Island Research Station, with particular appreciation to Ainsley and Paul Carin, Anne Hoggett, Somei Jonda, Peter and Jane Miller, and Lyle Vail for their assistance with our field trips. We would like to thank staff at the Sydney Institute of Marine Science for help while carrying out laboratory work in the facility.

