## Supplementary Material for "Population genomics highlight the vulnerability of coral-dwelling gobies to ecological losses due to climatic disturbances"

Supplementary Material for Manuscript Titled:

TBA

Suppl. Tab. 1. Amova results showing the difference among three locations for *Gobiodon* species. Samples sizes included as denoted by n. Number of variable loci that passed filtering parameters denoted by no. loci. PNG = Papua New Guinea, LI = Lizard Island, OTI = One Tree Island.

| | Source of Variation | d.f. | SS | %Variation | F-value | P |
|---|---|---|---|---|---|---|
| *Gobiodon fuscoruber* | | | | | | |
| PNG: n = 36<br>LI: n = 34 | Among locations | 2 | 1720.362 | 14.44 | 0.14435 | <0.0001 |
| OTI: n = 31<br>no. loci = 5751 | Within locations | 199 | 13873.262 | 85.56 | | |
| *Gobiodon histrio* | | | | | | |
| PNG: n = 36<br>LI: n = 39 | Among locations | 1 | 586.881 | 2.90 | 0.02896 | <0.0001 |
| no. loci = 10006 | Within locations | 148 | 26862.399 | 97.10 | | |
| *Gobiodon quinquestrigatus* | | | | | | |
| PNG: n = 41<br>LI: n = 29 | Among locations | 2 | 712.423 | 5.00 | 0.05003 | <0.0001 |
| OTI: n = 29<br>no. loci = 6527 | Within locations | 195 | 15697.294 | 95.00 | | |
| *Gobiodon rivulatus* | | | | | | |
| PNG: n = 30<br>LI: n = 45 | Among locations | 2 | 1121.252 | 8.92 | 0.08916 | <0.0001 |
| OTI: n = 28<br>no. loci = 6345 | Within locations | 203 | 15058.525 | 91.08 | | |

Suppl. Tab. 2. Pairwise $F_{ST}$ values and corresponding p-values between locations of *Gobiodon fuscoruber* calculated through Arlequin software. Below the grey line are the pairwise $F_{ST}$ values between the given locations, while above the grey line are the corresponding p-values for the $F_{ST}$ statistic. PNG = Papua New Guinea, LI = Lizard Island, OTI = One Tree Island.

| | PNG | LI | OTI |
|---|---|---|---|
| PNG | | <0.0001 | <0.0001 |
| LI | 0.11612 | | <0.0001 |
| OTI | 0.22041 | 0.10689 | |

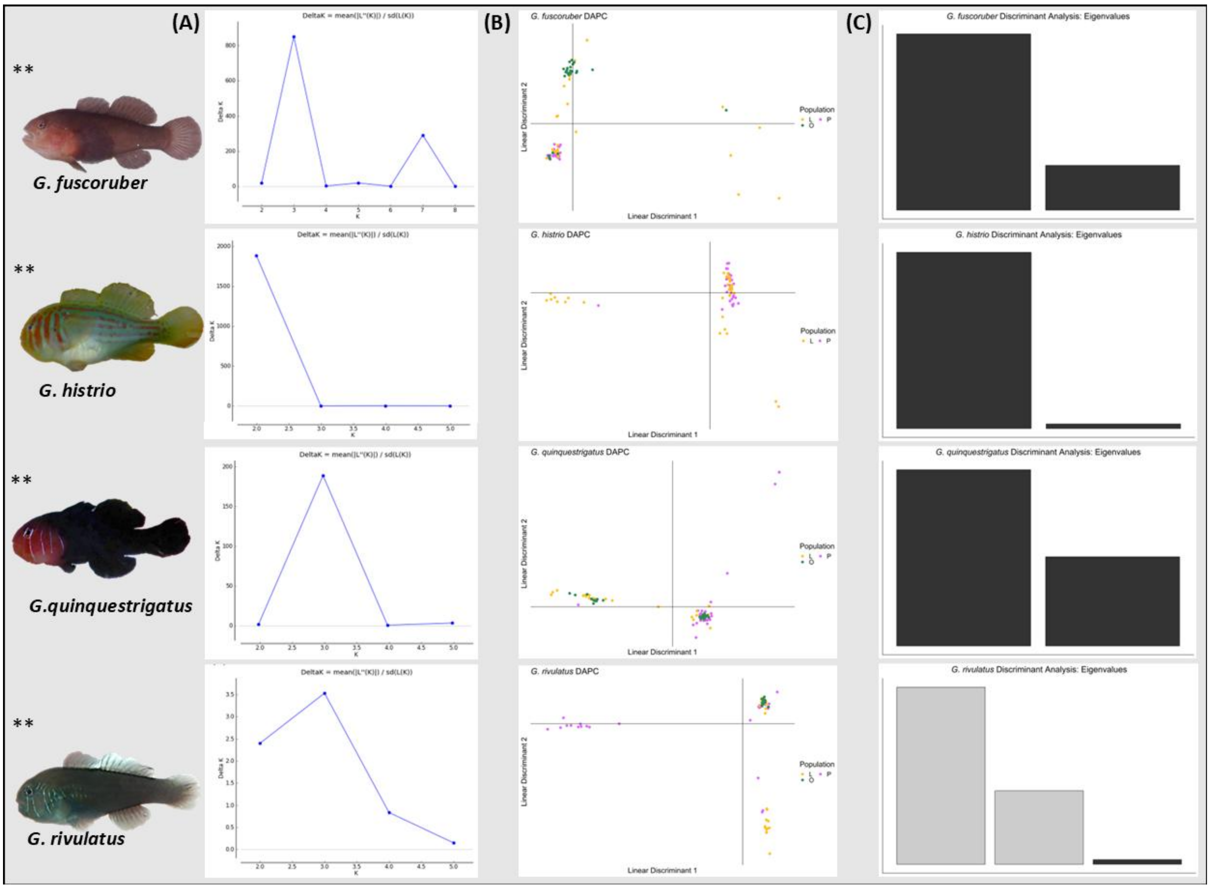


Suppl. Fig. 1. (a) DeltaK graphs from HARVESTER determining the best K-value, (b) Discriminant Analysis Principal Components and (c) eigen values for each axis for species compared among locations (purple = Papua New Guinea, orange = Lizard Island, Green = One Tree Island); asterisks mean significant differences observed among groups.

Suppl. Tab. 3. Pairwise $F_{ST}$ values and corresponding p-values between sampled locations of *Gobiodon quinquestrigatus* calculated through Arlequin software. Below the grey line are the pairwise $F_{ST}$ values between the given populations, while above the grey line are the corresponding p-values for the $F_{ST}$ statistic.

| | PNG | LI | OTI |
|---|---|---|---|
| PNG | | <0.0001 | <0.0001 |
| LI | 0.06381 | | <0.0001 |
| OTI | 0.04996 | 0.02850 | |

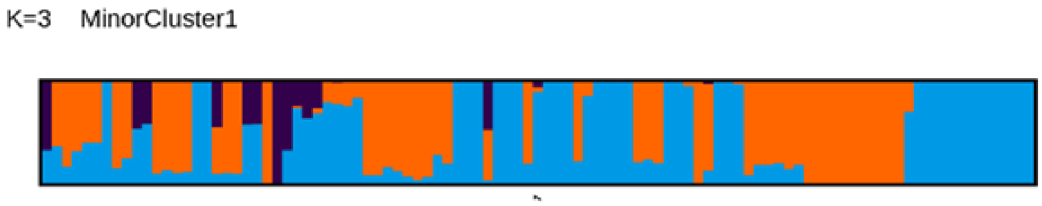


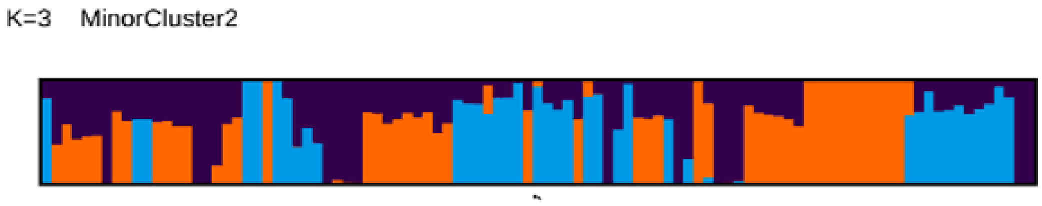


Suppl. Fig. 2. Two minor modes were created for *Gobiodon quinquestrigatus* with a K-value of three.

Suppl. Tab. 4. Pairwise $F_{ST}$ values and corresponding p-values between locations of *Gobiodon rivulatus* calculated through Arlequin software. Below the grey line are the pairwise $F_{ST}$ values between the given locations, while above the grey line are the corresponding p-values for the $F_{ST}$ statistic.

| | PNG | LI | OTI |
|---|---|---|---|
| PNG | | <0.0001 | <0.0001 |
| LI | 0.09991 | | <0.0001 |
| OTI | 0.12599 | 0.04345 | |

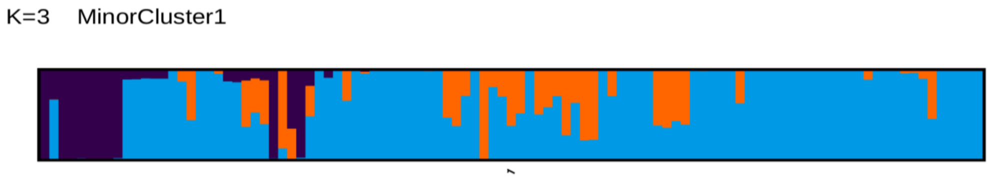


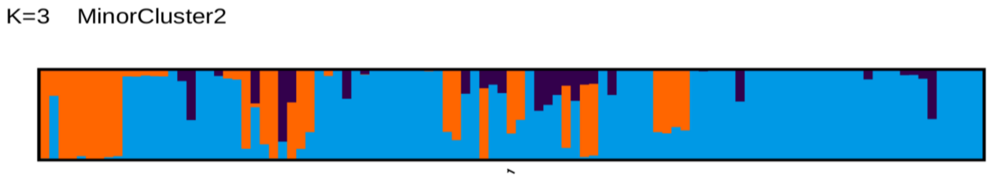


Suppl. Fig. 3. Two minor modes were created for *Gobiodon rivulatus* with a K-value of three.

Suppl. Tab. 5. Amova results of pre- and post-disturbances at Lizard Island for *Gobiodon* species. Number of variable loci that passed filtering parameters denoted by no. loci.

| | Source of Variation | d.f. | SS | %Variation | F-value | P |
|---|---|---|---|---|---|---|
| *Gobiodon brochus* | | | | | | |
| PRE: n = 31<br>POST: n = 31 | Among groups | 1 | 361.895 | 0.07 | 0.00073 | 0.87031 |
| no. loci = 9724 | Within groups | 122 | 42233.952 | 99.93 | | |
| *Gobiodon erythrospilus* | | | | | | |
| PRE: n = 31<br>POST: n = 31 | Among groups | 1 | 391.081 | 0.53 | 0.00529 | 0.01751 |
| no. loci = 10760 | Within groups | 122 | 35882.387 | 99.47 | | |
| *Gobiodon histrio* | | | | | | |
| PRE: n = 39<br>POST: n = 31 | Among groups | 1 | 1336.583 | 7.86 | 0.07859 | <0.0001 |
| no. loci = 9391 | Within groups | 138 | 26760.959 | 193.92000 | | |
| *Gobiodon quinquestrigatus* | | | | | | |
| PRE: n = 29<br>POST: n = 30 | Among groups | 1 | 859.483 | 7.88 | 0.07885 | <0.0001 |
| no. loci = 7656 | Within groups | 116 | 16482.440 | 92.12 | | |

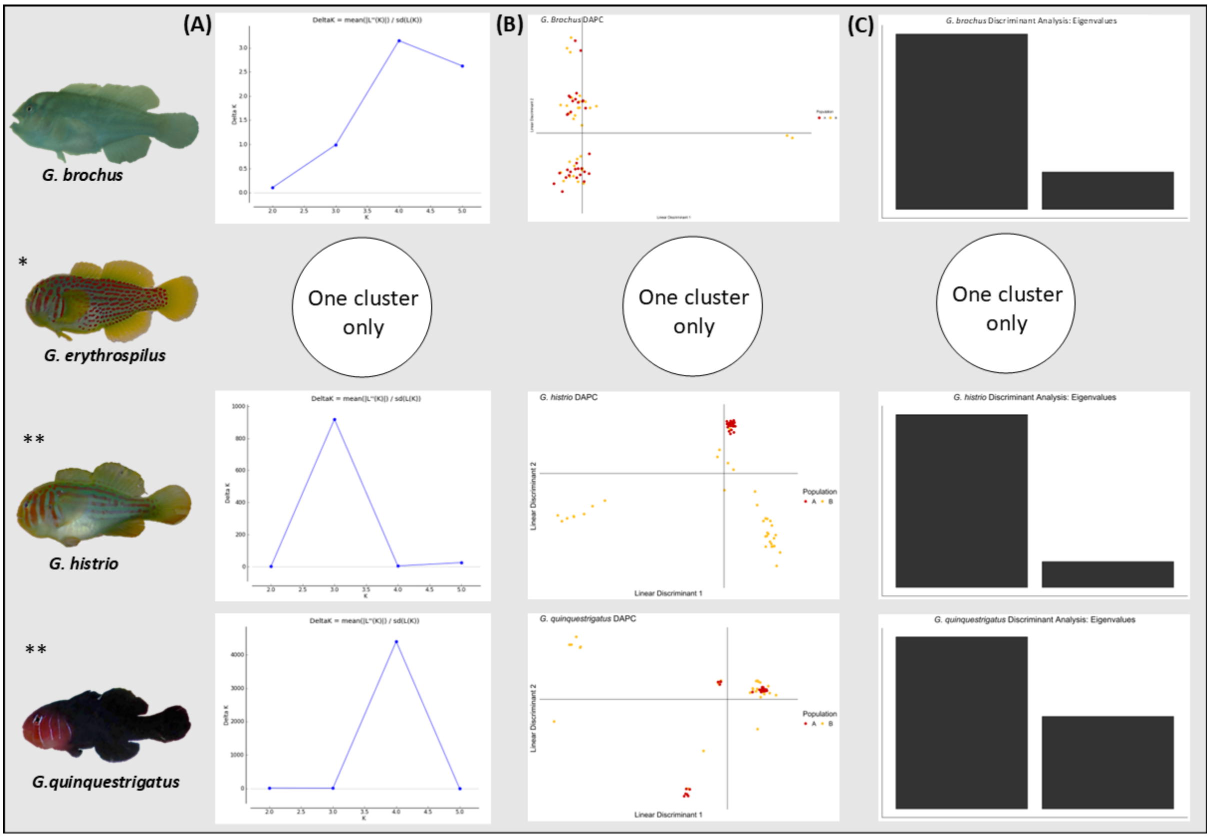


Suppl. Fig. 4. (a) DeltaK graphs from HARVESTER determining the best K-value, (b) Discriminant Analysis Principal Components and (c) eigen values for each axis for species compared pre-disturbances (orange) and post-disturbances (dark red); ** means $p < 0.001$; * means $p < 0.02$.
